# Single-cell RNA-seq of human induced pluripotent stem cells reveals cellular heterogeneity and cell state transitions between subpopulations

**DOI:** 10.1101/119255

**Authors:** Quan H. Nguyen, Samuel W. Lukowski, Han Sheng Chiu, Anne Senabouth, Timothy J. C. Bruxner, Angelika N. Christ, Nathan J. Palpant, Joseph E. Powell

## Abstract

Heterogeneity of cell states represented in pluripotent cultures have not been described at the transcriptional level. Since gene expression is highly heterogeneous between cells, single-cell RNA sequencing can be used to identify how individual pluripotent cells function. Here, we present results from the analysis of single-cell RNA sequencing data from 18,787 individual WTC CRISPRi human induced pluripotent stem cells. We developed an unsupervised clustering method, and through this identified four subpopulations distinguishable on the basis of their pluripotent state including: a core pluripotent population (48.3%), proliferative (47.8%), early-primed for differentiation (2.8%) and late-primed for differentiation (1.1%). For each subpopulation we were able to identify the genes and pathways that define differences in pluripotent cell states. Our method identified four transcriptionally distinct predictor gene sets comprised of 165 unique genes that denote the specific pluripotency states; and using these sets, we developed a multigenic machine learning prediction method to accurately classify single cells into each of the subpopulations. Compared against a set of established pluripotency markers, our method increases prediction accuracy by 10%, specificity by 20%, and explains a substantially larger proportion of deviance (up to 3-fold) from the prediction model. Finally, we developed an innovative method to predict cells transitioning between subpopulations, and support our conclusions with results from two orthogonal pseudotime trajectory methods.

## Introduction

The transcriptome is a key determinant of the phenotype of a cell and regulates the identity and fate of individual cells. Much of what we know about the structure and function of the transcriptome comes from studies averaging measurements over large populations of cells, many of which are functionally heterogeneous. Such studies conceal the variability between cells and so prevent us from determining the nature of heterogeneity at the molecular level as a basis for understanding biological complexity. Cell-to-cell differences in any tissue or cell culture are a critical feature of their biological state and function.

In recent decades, the isolation of pluripotent stem cells, first in mouse followed by human (Evans and Kaufman, 1981; Thomson et al., 1998), and the more recent discovery of deriving pluripotent stem cells from somatic cell types (iPSCs) (Takahashi and Yamanaka, 2006), is a means to study lineage-specific mechanisms underlying development and disease to broaden our capacity for biological therapeutics (Palpant et al., 2017). Pluripotent stem cells are capable of unlimited self-renewal and can give rise to specialized cell types based on stepwise changes in the transcriptional networks that orchestrate complex fate choices from pluripotency into differentiated states.

In addition to individual published data, international consortia are banking human induced pluripotent stem cells (hiPSCs) and human embryonic stem cells (hESCs) and providing extensive phenotypic characterization of cell lines including transcriptional profiling, genome sequencing, and epigenetic analysis as data resources (Streeter et al., 2017; The Steering Committee of the International Stem Cell, 2005). These data provide a valuable reference point for functional genomics studies but continue to lack key insights into the heterogeneity of cell states that represent pluripotency.

While transcriptional profiling has been a common endpoint for analyzing pluripotency, the heterogeneity of cell states represented in pluripotent cultures has not been described at a global transcriptional level. Since each cell has a unique expression state comprising a collection of regulatory factors and target gene behavior, single-cell RNA sequencing (scRNA-seq) can provide a transcriptome-level understanding of how individual cells function in pluripotency (Wen and Tang, 2016). These data can also reveal insights into the intrinsic transcriptional heterogeneity comprising the pluripotent state. In this study, we provide the largest dataset of single-cell transcriptional profiling of undifferentiated hiPSCs currently available, which cumulatively amount to 18,787 cells across five biological replicates. Moreover, we have developed several innovative single-cell methods focused on unbiased clustering, machine-learning classification, and quantitative and directional cellular trajectory analysis. Our findings address the following hypotheses: (1) pluripotent cells form distinct groups or subpopulations of cells based on biological processes or differentiation potential; (2) transcriptional data at single cell resolution reveal gene networks governing specific cell subpopulations; (3) transcripts can exhibit differences in gene expression heterogeneity between specific subpopulation of cells.

## Results

### Description of the parental hiPSC line, CRISPRi

WTC-CRISPRi hiPSCs (Mandegar et al., 2016) were chosen as the parental cell line for this study. These cells are genetically engineered with an inducible nuclease-dead Cas9 fused to a KRAB repression domain (**Supplemental Figure S1a**). Transcriptional inhibition by gRNAs targeted to the transcriptional start site is doxycycline-dependent and can be designed to silence genes in an allele-specific manner. The versatility of this line provides a means to use this scRNA-seq data as a parental reference point for future studies aiming to assess the transcriptional basis of pluripotency at the single cell level. Cells were verified to have a normal 46 X,Y male karyotype by Giemsa banding analysis before analysis by scRNA-seq (**Supplemental Figure S1b**).

### Overview of single-cell RNA sequence data

After quality control of the sequencing data (Methods), we obtained 1,030,909,022 sequence reads for 20,482 cells from five hiPSC single cell samples (**Supplemental Table S1, Supplemental Figure S2**), with 63-71% confidently and uniquely mapped (mapping quality 255) to the human reference transcriptome hg19 (Ensembl, release 75). We sequenced 19,937 cells from four samples to an average depth of 44,506 reads-per-cell (rpc), and one sample consisting of 545 cells was sequenced to an average depth of 318,909 rpc. On average, 2,536 genes and 9,030 Unique Molecular Identifiers (UMIs) were detected per cell. Comparing the sequencing results between five samples, we observe little variation in the total number of genes detected per sample, despite differences in the number of cells per sample, and average reads per cell (**Supplemental Table S1**). For example, we observed only a slight increase in the average number of genes detected for cells sequenced at a greater depth (**Supplemental Tables S1, S3, Supplemental Figure S2f**), and no gain in the total number of genes detected for all cells in the whole sample. These results suggest that an average of 44,506 rpc achieves close to maximum total gene detection in our samples. We found that the number of reads per cell affects per-cell gene detection (sensitivity), whereas the number of cells per sample effects on total gene detection (more unique genes per sample). Overall, after quality control we detected 16,064 unique genes, which were expressed in at least 1% of the total cells. Importantly, of the 16,064 genes, only one was unique to a single sample (**Supplemental Table S4**). We subsequently removed 1,738 cells due to a high percentage of expressed mitochondrial and/or ribosomal genes (**Methods, Supplemental Table S2**), leaving a total of 18,787 high-quality hiPSCs for further analysis. Following between-sample and between-cell normalisation, we observed no evidence for batch effects due to sample or sequencing run (**Figure 1a, Supplemental Figure S3**).

**Figure 1.**
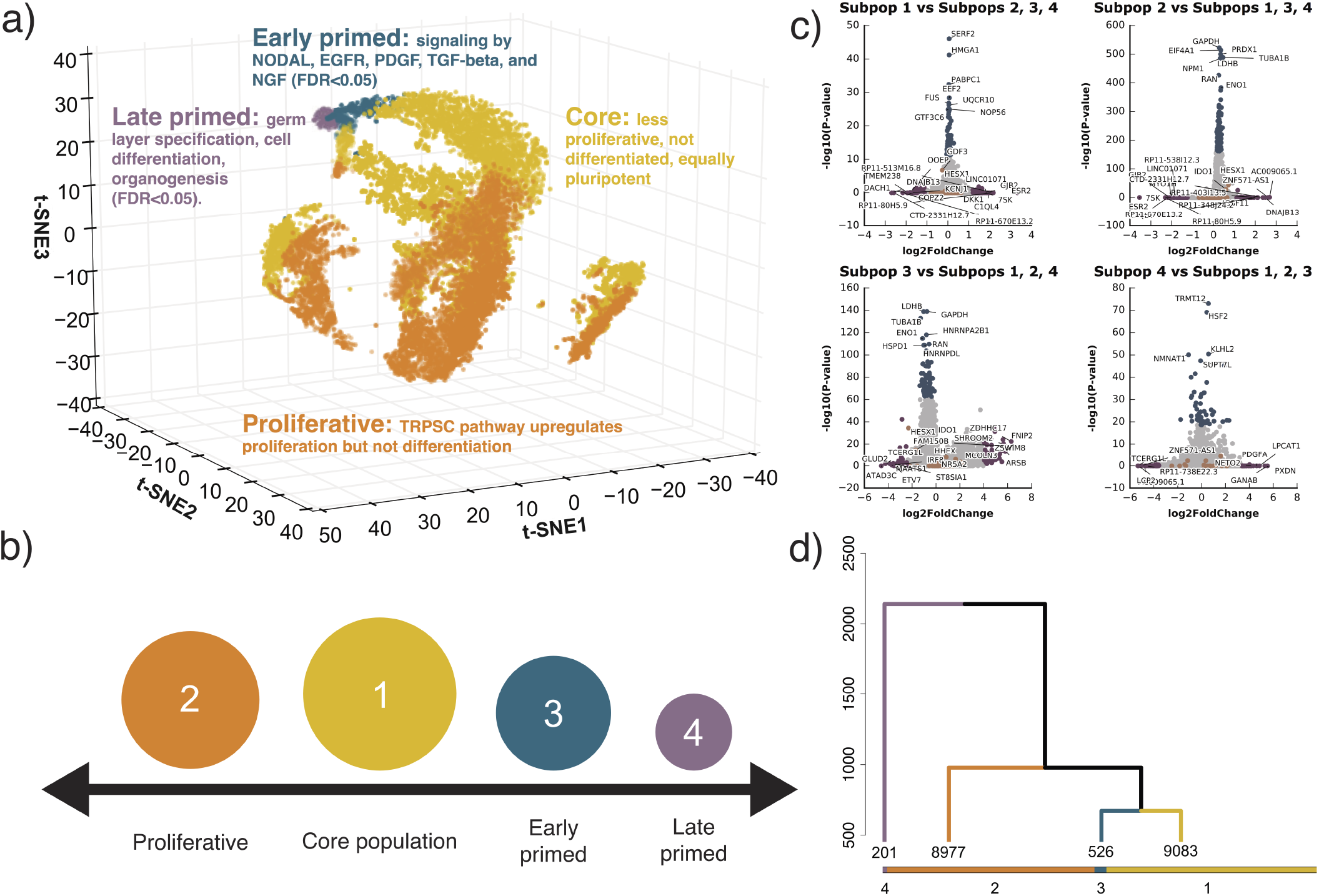
Identification of four cell subpopulations from 18,787 hiPSC cells, sequenced from five biological replicates. (**a**) Three-dimensional t-SNE distribution of cells based on gene expression value. Each point represent a single cell in three-dimensional space. A t-SNE transformation of the data was used for positioning cells, while four cell subpopulation labels (marked by different colors) represent results from clustering, and are independent of t-SNE data transformation (see http://computationalgenomics.com.au/shiny/hipsc/ for interactive, searchable figure). Pathway analysis based on differential expression identified functional properties that distinguish each subpopulation. (**b**) Four pluripotent subpopulations functionally separated from a homogeneous hiPSC population. (**c**) The top significantly differentially expressed genes of cells in a subpopulation compared to cells in the remaining three subpopulations. Genes denoted with orange points are known naive and primed markers. Genes represented with blue and purple points are those in the top 0.5% highest logFC or −log(*P-value*) respectively. (**d**) Unsupervised clustering of all cells into four subpopulations. The dendrogram tree displays distance and agglomerative clustering of the cells. Each branch represents one subpopulation. The clustering is based on a dynamic tree cut that performs a bottom-up merging of similar branches. The number of cells in each of the four subpopulations are given below branches.

### Identification of four hiPSC subpopulations based on biological function

We developed an innovative classification method, which we term Unsupervised High-Resolution Clustering (UHRC), to objectively assign cells into subpopulations based on genome-wide transcript levels (**Figure 1, Supplemental Figure S4**, and **Methods** section). The UHRC procedure is comprised of three unbiased algorithms. First, a PCA reduction step was implemented to overcome the inherent multicollinearity in single cell expression data. Subsequently we apply bottom-up agglomerative hierarchical clustering, which importantly provides a ‘data-driven’ identification of clusters, rather than inputting a specific number of expected clusters, as is the case with K-means algorithms. Third, to robustly define large clusters, a dynamic branch merging process is used to detect complex nested structures and outliers. This unbiased method identified four independent subpopulations of cells containing 48.3, 47.8, 2.8 and 1.1 percent of the 18,787 cells, respectively (**Figure 1 a, b and d**; and **Supplemental Figure S4**). Importantly, after clustering, we did not observe evidence for batch effects underlying any of the four cell subpopulations (**Figure 1a, Supplemental Table S5**), suggesting that the clusters represent biological and not technical factors. By comparing gene expression levels between subpopulations, we identified four differentially expressed gene sets that distinguish each subpopulation from the remaining cells (**Figure 1c, Supplemental Table S6**). We have made this data available via an interactive, gene-searchable web application available at http://computationalgenomics.com.au/shiny/hipsc/.

We initially examined transcript dynamics in the subpopulations based on expression of previously described markers of pluripotency and lineage determination (Tsankov et al., 2015) (**Figure 2** and **Supplemental Table S7**). Of the 18,787 cells, 99.8% expressed at least one of 19 pluripotency genes (**Supplemental Table S8**). Furthermore, genes with known roles in pluripotency had stronger expression across all subpopulations compared to genes involved in lineage determination (**Figure 2a-b, Supplemental Tables S7 and S8**). For example, *POU5F1* (also known as *OCT4*), which encodes a transcription factor critically involved in the self-renewal of undifferentiated pluripotent stem cells (Boyer et al., 2005), was consistently expressed in 98.6% of cells comprising all four subpopulations (**Figure 2a-b, Supplemental Tables S7 and S8**). Other known markers of pluripotency such as *SOX2, NANOG* and *UTF1* were expressed across the subpopulations (**Figure 2a-b, Supplemental Tables S7 and S8**), but showed differences in expression heterogeneity, suggesting differences in the pluripotent state across subpopulations (**Supplemental Table S7**).

**Figure 2.**
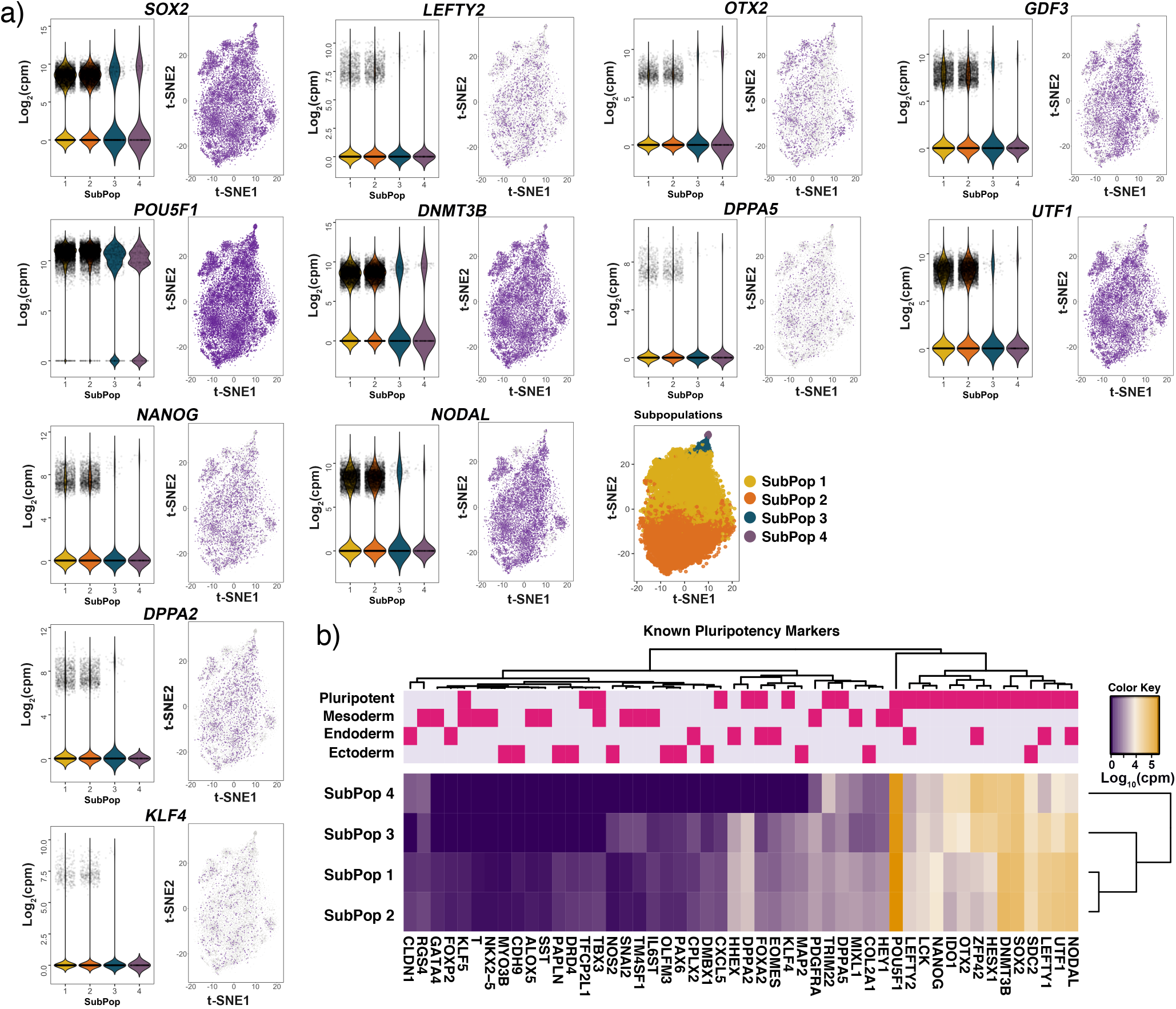
Expression levels of known pluripotency and lineage primed markers. (**a**) Violin and jitter plots for expression of top pluripotency markers and expression of the selected genes represented by t-SNE plots. Each point represents a single cell. Color gradient in the t-SNE plot represents relative expression level of the gene in a cell across the whole population and subpopulations (light grey = low; dark purple = high). (**b**) Heatmap of the mean expression of known markers within each subpopulation. The upper panel shows the classifications of genes into pluripotency and lineage-primed markers.

### Classification of hiPSC subpopulations

We sought to identify biological processes underlying transcriptional classification of cell subpopulations by firstly performing a statistical analysis to identify significantly differentially expressed genes between subpopulations (**Methods, Figure 1c, Supplemental Table S6**). Differentially expressed genes with a fold-change significant at a Bonferroni-corrected p-value threshold (*p* < 3.1 × 10^−7^) were evaluated for enrichment of functional pathways (**Supplemental Tables S9-S13**).

Cells classified in subpopulations one and two, which comprise 96.1% of total cells analyzed (**Figure 1 a, b, c**), were distinguished from one another by significantly different expression levels of genes in alternate pathways controlling pluripotency and differentiation (**Supplemental Figure S5, Supplemental Tables S9-S11**). The Transcriptional Regulation of Pluripotent Stem Cells (TRPSC) pathway was consistently up-regulated in cells classified as subpopulation two compared to subpopulation one (**Supplemental Figure S5, Supplemental Table S11 and S14**). TRPSC is an auto-activation loop, which maintains expression of *POU5F1, NANOG*, and *SOX2* at high levels. Complexes containing various combinations of these transcription factors (Lam et al., 2012) can activate the expression of genes whose products are associated with rapid cell proliferation, and also repress the expression of genes associated with cell differentiation (Forristal et al., 2010; Guenther, 2011) (**Supplemental Figure S5**). In particular, *POU5F1, NANOG*, and *SOX2* are more highly expressed in subpopulation two (**Supplemental Table S7**), and the direction of differential expression of genes associated with cell proliferation and repression of cell differentiation (Forristal et al., 2010; Guenther, 2011) is consistent with subpopulation two containing cells that are more active in their self-renewal than cells in subpopulation one (**Supplemental Tables S7, S10 and S14**).

We hypothesized that expression differences between subpopulations one and two could reveal novel, subpopulation-specific markers. We applied unbiased differential expression analysis between subpopulation one and two, identifying 49 statistically significant genes that were taken forward to biological pathway analysis (**Supplemental Table S6, Supplemental Figure S5**). Among these 49 genes, two key transcription factors and a signaling receptor were significantly higher in subpopulation two than subpopulation one, including *SALL4* (Spalt-like transcription factor 4, *p-adjusted:* 7.0 × 10^−5^), *ZIC1* (Zic family member 1, *p-adjusted*: 4.3×10^−5^), and *NR6A1* (Nuclear Receptor Subfamily 6 Group A Member 1, *p-adjusted*: 3.7 × 10^−6^) (**Supplemental Table S14**).

*SALL4* is one of the key transcription factors that participates in controlling transcriptional balance in pluripotent cells and suppressing differentiation (Miller et al., 2016) (**Supplemental Figure S5**). Specifically, *SALL4* activates transcription of *POU5F1* and maintains pluripotency (Yang et al., 2010b). *ZIC1*, upregulated gene in subpopulation two, was identified by GeneMANIA analysis to be related to *SALL4* through shared protein domains (**Supplemental Figure S6**). Both *ZIC1* and *SALL4* were predicted by the STRING database to interact with key pluripotency markers (**Supplemental Figure S6**). Furthermore, *ZIC1* and its paralog *ZIC3*, a key member in the *TRPSC* pathway (**Supplemental Figure S5**), are involved in maintaining the undifferentiated state, for example in the case of neural precursor cells (Inoue et al., 2007). Moreover, we also identified another differentially expressed gene, *NR6A1*, a known regulator of *POU5F1* later in differentiation (Fuhrmann *et al*. 2001; Weikum *et al*. 2017), and which we predict is likely to participate in the *TRPSC* pathway since its paralog, *NR5A1*, is among the key members of this pathway (Supplemental Figure S6). Based on these observations, we hypothesize that in subpopulation two, three differentially expressed (DE) genes, *SALL4, NR6A1* and *ZIC3*, cooperate with key pluripotency transcription factors *POU5F1, SOX2*, and *NANOG* to activate genes related to proliferation, but not genes involved in differentiation (Supplemental Figure S5).

Compared to subpopulations one and two, subpopulations three and four represent pluripotent populations with significant down-regulation of key pluripotency network genes (e.g. *NANOG* and *UTF1)* (**Figure 2a-b**). For all known pluripotency markers examined in **Supplemental Table S8**, we observed expression in a higher proportion of cells in subpopulation one and two than subpopulations three and four. Furthermore, the mean expression of 46 out of 56 known markers involved in pluripotency shown in **Supplemental Table S7** is higher in subpopulations one and two than in three and four. For subpopulation three, comprising 2.8% of cells, Reactome pathway enrichment analysis of 2,534 DE genes between subpopulations three and four showed the top pathways related to developmental signaling and transcriptional regulation via chromatin modification (**Supplemental Table S12**). Intracellular signaling pathways that control cell proliferation, cell differentiation, and cell migration, such as *EGFR, PDGF*, and *NGF* pathways (FDR < 1.7 × 10^−6^), were the top three most enriched pathways (**Supplemental Table S12**). Additionally, signaling pathways by *FGFRs* involved in differentiation were also significantly enriched (FDR < 3 × 10^−4^). Comparing subpopulations three and one, signaling by *TGF*-beta, and signaling by *NODAL* were in the top enriched pathways (FDR < 8 × 10^−3^). Similarly, signaling by *NODAL* (FDR < 0.04) (Levin cent *et al*., 2003) and pre-NOTCH processing (FDR < 0.04) (Artavanis-Tsakonas et al., 1999), which are involved in cell fate decisions, were in the top enriched pathways when comparing subpopulation three to subpopulations one and two (**Supplemental Table S12**). In addition, ‘chromatin modifying enzymes’ is among the top Reactome pathways distinguishing subpopulation three, suggesting that cells are undergoing active transcription regulation. Thus, pluripotent cells in subpopulation three appear more lineage primed compared to subpopulations one and two.

Further, we extended the pathway enrichment analysis to a broad collection of background gene ontology databases, beyond the manually curated Reactome database, by using BiNGO in Cytoscape for all 1,706 DE genes in subpopulation four vs. all other subpopulations (1.1% of analyzed cells) (**Supplemental Table S13**). We found the top enriched pathways related to differentiation including genes involved in: gastrulation (FDR < 1.3 × 10^−2^) and formation of primary germ layer (FDR < 1.4 × 10^−2^); developmental process (FDR < 2.8 × 10^−3^); and cell differentiation (FDR < 1.2 × 10^−2^); and more than 20 significantly enriched pathways related to organogenesis (FDR < 5 × 10^−2^; Supplemental Table S13). Thus, although cells in subpopulation four are still pluripotent, as indicated by the expression of pluripotent markers, they likely represent cells at a late-primed state progressing toward differentiation.

Taken together, our transcriptional profiling of single cells revealed four subpopulations defined by their pluripotency levels, cell proliferation, and potential for cell lineage commitment. Subpopulation one pluripotent cells likely represent a core pluripotent state, subpopulation two proliferative pluripotent cells, subpopulation three as early-primed for differentiation, and subpopulation four as late-primed for differentiation (**Figure 1b**).

### Improved cell classification using a reduced set of gene markers

Current methods of classifying cells into sub-types or functional groups rely on prior knowledge of so-called ‘marker genes’ to inform the pseudotime algorithm. One limitation of this approach is that, to discover new or rare cell types using single-cell RNA-seq, the use of established markers as an input may not be feasible.

Using differentially expressed genes identified between subpopulations, we built a novel unbiased machine learning predictor to identify the pluripotency potential of a single cell. To avoid over-fitting the model due to co-expression of genes, we used a variable selection regression model called LASSO (Tibshirani, 1996) to estimate gene effects differentiating each subpopulation conditional upon the effects of other genes. Using a 100-fold bootstrapping approach, we estimated the predictive accuracy of identifying a cell in each of the four subpopulations (**Figure 3, Supplemental Table S15**) (Tibshirani, 1996). To detect new gene markers compared to the use of known pluripotency markers (**Supplemental Table S7**), we applied LASSO to selected sets of differentially expressed genes between one subpopulation compared to the remaining subpopulations. Consistently across four comparisons, our method increases prediction accuracy by 10%, specificity by 20%, and explains a substantially larger proportion of deviance (up to 3-fold) from the prediction model compared to known markers (**Supplemental Figures S7-S8, Supplemental Tables S15-S16**). Prediction by DE genes showed markedly higher performance, measured specifically by sensitivity, specificity and percentage of deviance explained, in distinguishing cells classified in subpopulations three and four from the remaining cells (**Figure 3a**). Similarly, a significantly higher percent of deviance can be explained by using DE genes compared to using known markers (*t*-test, *p*=1.4×10^−77^) (**Figure 3b**). We observed the highest classification accuracy using genes identified using the LASSO model for cells in subpopulations three and four than cells in subpopulations one and two, suggesting that these subpopulations were more divergent from the remaining majority of the cell population (**Figure 3c**). This observation further supports the classification of subpopulations three and four as more primed to differentiation than subpopulations one and two. Although the difference in accuracy was less apparent for cells in subpopulations one and two, the improved accuracy achieved using the DE genes compared to known markers supports the conclusion that our analysis has identified novel genes that act as pluripotency markers (**Figure 3c**).

**Figure 3.**
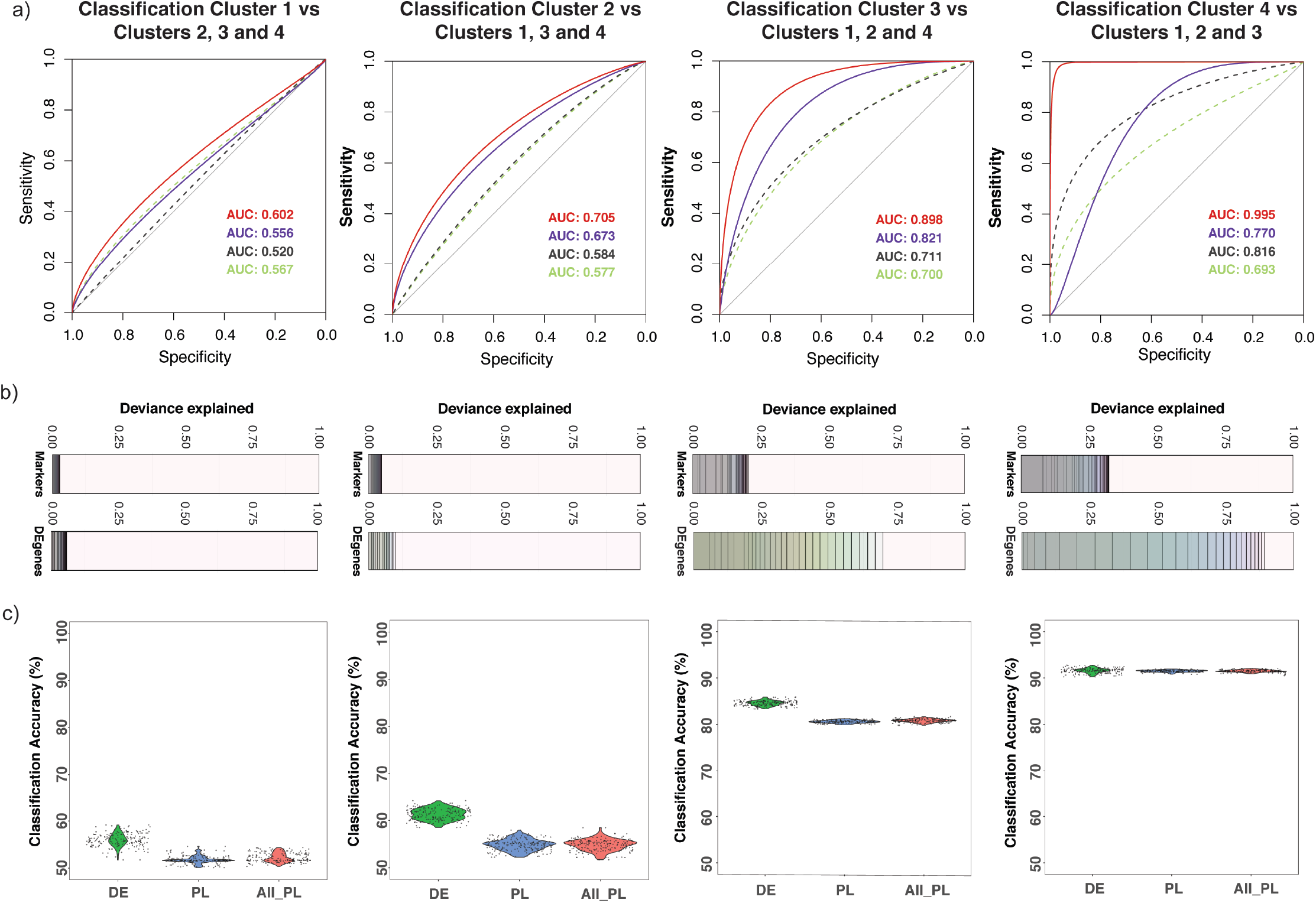
Selection of significant gene predictors for classifying each subpopulation using LASSO regression. (**a**) For each subpopulation, a LASSO model was run using a set of differentially expressed (DE) genes and another set of known markers. Dashed lines are ROC (Receiver Operating Characteristic) curves for models using known markers. Continuous lines are for models using differentially expressed genes. The text shows corresponding AUC (Area Under the Curve) values for ROC curves. For each case (known markers or DE genes), a model with the lowest AUC and another model with the highest AUC are given. Lower AUC values (and ROC curves) in the prediction models using known markers suggested that the models using DE genes performed better in sensitivity and specificity. (**b**) Each deviance plot (bottom panel) shows the deviance explained (x-axis) by a set of gene predictors (numbers of genes is shown as vertical lines and varies from 1 to maximum value as the total number of gene input or to the minimum number of genes that can explain most of the deviance). The remaining space between the last gene and 1.0 border represents deviance not explained by the genes in the model. (**c**) Classification accuracy calculated using a bootstrap method using all known markers (both pluripotent markers and primed lineage markers) or markers from our differentially expressed gene list is shown. Expression of LASSO selected genes for subpopulation one and subpopulation two is shown in Supplemental Figure S7. The X-axis labels are for three cases: using LASSO selected differentially expressed genes (DE); LASSO selected pluripotency/lineage-primed markers (PL); and all pluripotency/lineage-primed markers (All PL).

### Cell transition between pluripotency states

We next sought to investigate cell trajectories between subpopulations of iPSCs. We further deployed the results from the LASSO variable selection procedure described above to develop a classifier model, termed Local Transitions between Subpopulations (LTS) that can predict the conversion potential of cells in one subpopulation into cells in a target subpopulation. For each subpopulation, we trained the expression data using the genes identified as differentially expressed, to optimize a logistic regression model by penalized maximum likelihood estimation. The training provides an optimal LASSO model with selected genes and corresponding coefficients. The coefficients estimated from the training model were fitted into the expression levels of each of the remaining subpopulations (target populations), enabling us to predict which cells in the target population were the most similar to cells in the training subpopulation. The percentage of cells within the target subpopulation accurately predicted by the training subpopulation is termed the transition score. The transition scores provide both a quantitative and directional estimate of the progression potential between subpopulations. Using LTS to predict lineage trajectory between subpopulations of pluripotent cells showed that subpopulation one is capable of progressing into all other states, with the potential to convert into approximately 45%, 69%, and 47% of the cells in the subpopulation two, three, and four respectively (**Table 1, Figure 4e**). In contrast, subpopulation two predominantly transitions into subpopulation one (39%), with low transition scores when compared against subpopulations three and four (< 5%), (**Figure 4e**). The difference in transition scores from subpopulation two to other subpopulations suggests that the cells were at a reversible, self-renewal state (**Table 1**). Notably, the LASSO method predicts the highest transition score for the progression from subpopulation three to subpopulation four (> 99%), providing further support that subpopulation three is “early primed” prior to the “late primed” state in subpopulation four (**Figure 4e**).

**Figure 4.**
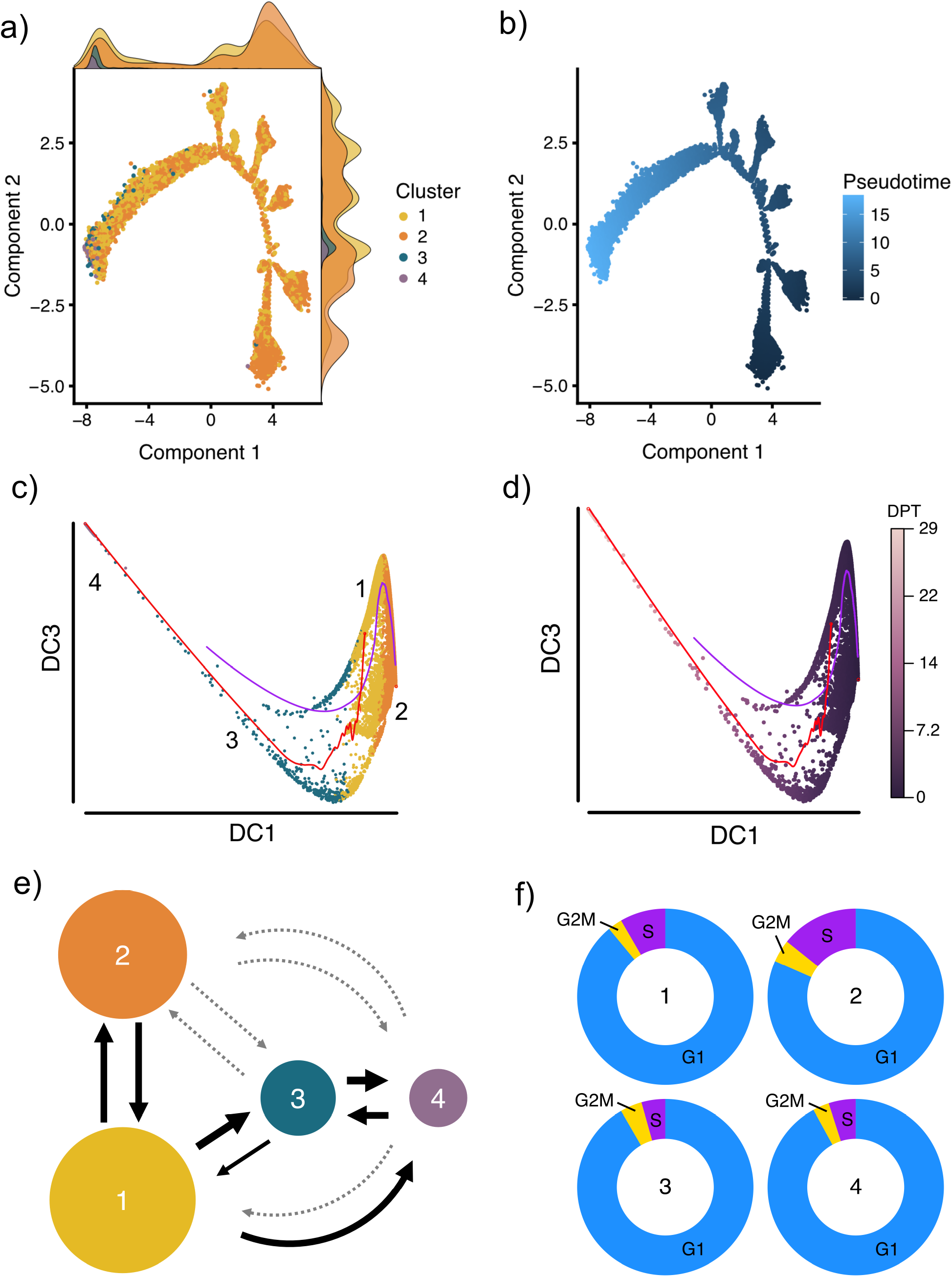
Trajectory and cell cycle membership analysis. Cell differentiation potential was mapped using two pseudotime approaches implemented in Monocle2 2 and Destiny and a novel transition estimation method. Panel (**a**) shows the results of the Monocle2 analysis, colored by subpopulation, and the normalized density of the cells in each location along the trajectory is shown as a density curve in the *x* and *y* plot margins. Panel (**b**) shows the differentiation distance from the root cell to the terminal state where the dark blue represents the beginning (the root) and light blue represents the end (the most distant cells from the root) of the pseudotime differentiation pathway. (**c**) and (**d**) show the results of diffusion pseudotime analysis, colored by cluster (**c**) and by diffusion pseudotime (**d**). DC refers to diffusion component and DPT refers to diffusion pseudotime. The red and blue pathways in c and d represent the transition path from cell to cell calculated by a random-walk algorithm. (**e**) We developed a novel approach that uses the LASSO classifier to quantify directional transitions between subpopulations. Panel (**e**) shows the percent of transitioning cells predicted between subpopulations. The weight of the arrows is relative to percentage (thicker is higher percentage), and the light grey dotted arrows represent percentages lower than 20. Cell cycle stages were predicted for each cell by subpopulation (**f**). Subpopulation 1 (‘Core’) contains a significantly lower number of cells in the S phase (synthesis) compared to Subpopulation 2 (‘Proliferative’; Fisher’s exact Test, P < 2.2×10−16).

**Table 1.** Percentage of cells in other subpopulations predicted by LASSO

|  | Subpop 1 | Subpop 2 | Subpop 3 | Subpop 4 |
| --- | --- | --- | --- | --- |
| Subpop 1 | - | 45.1 | 69.4 | 47.3 |
| Subpop 2 | 39.4 | - | 4.9 | 2.5 |
| Subpop 3 | 28.4 | 19.9 | - | 99.1 |
| Subpop 4 | 24.6 | 20.6 | 61.0 | - |

To confirm that the population of iPSCs was comprised of groups of cells located in different states along a differentiation path, we cross-validated our findings with two independent pseudotime analysis methods, namely Monocle2 (Qiu et al, 2017) and diffusion pseudotime (Haghverdi et al., 2016). These two methods estimate the differentiation distance of a cell compared to a root cell (see Methods section), but do not otherwise provide quantitative information about direction of movement. Monocle2 (**Figure 4a-b**) revealed a substantial overlap between subpopulations one and two, with cells classified in subpopulation one distributed across the trajectory, indicating a pluripotent state. Furthermore, the majority of cells within subpopulation two are located at the right terminal, indicating cells are closer to the root. As expected, cells within subpopulations three and four were located distal to cells from subpopulation two at the left terminal which supports our results suggesting classification of subpopulations three and four as being more primed to differentiate. The diffusion algorithm, implemented through the Destiny package (Angerer et al., 2016), maps cells onto a two-dimensional pseudotime space. The results obtained from diffusion pseudotime analysis (**Figure 4c-d**) were in strong agreement with the Monocle2 prediction, supporting our evidence that the four subpopulations present in a culture of iPSCs exist in multiple states of pluripotency.

### Validation against transcriptional data from the Human Induced Pluripotent Stem Cells Initiative (HipSci)

To confirm that genes selected by our LASSO analysis were also expressed in other hiPSC lines, we obtained open-access RNA-Seq transcript count data (tags per million - tpm) from the Human Induced Pluripotent Stem Cells Initiative (HipSci) for 71 hiPSC lines derived from reprogramming of dermal fibroblast biopsies from normal individuals (Streeter et al., 2017). Consistently, we observed expression of LASSO genes in 71 other independent hiPSC samples (**Supplemental Figure S8**). Moreover, we observed high correlation (*r* > 0.85) between the relative expression values among genes in our single-cell dataset with those genes in the HipSci bulk RNA-seq dataset (**Supplemental Figure S8c**). The high correlation further confirms that the single-cell sequencing data accurately reflects the relative abundance of transcripts.

### Transcriptional heterogeneity varies between cell subpopulations

With the large-scale data from 18,787 single cells greater than 16,000 genes were detected as expressed. Using this expression matrix, we were able to robustly analyze expression heterogeneity, both between genes, and for a given gene, between cells and cell subpopulations (**Supplemental Figure S9**). The inherently high heterogeneity of gene expression in scRNA-seq data, especially for low abundance genes with a more frequent on-off signal, may reduce the power to detect differential expression between cells (Shalek et al., 2013). Indeed, we identify more variation for subpopulations with smaller number of cells (**Supplemental Figure S9a**), and also for genes with low expression (**Supplemental Figure S9b**). Tagwise dispersion, which is expression variability for a gene across all cells in a subpopulation, decreased when average expression increased (**Supplemental Figure S9b**). The difference in the level of heterogeneity of gene expression for cells in a given subpopulation compared to other subpopulations is an important indicator of the relative dynamic cellular activity of the subpopulation. The red line in **Supplemental Figure S9b** shows the median dispersion of all genes across all cells within a subpopulation, thereby representing the average expression heterogeneity of the subpopulation. We found the median dispersion was higher in subpopulations three and four than in subpopulations one and two (**Supplemental Figure S9b**). This is consistent with our conclusion that subpopulations three and four were closer to a differentiated state compared to cells in subpopulations one and two, which are identified as being more pluripotent based on transcriptome analysis.

### Cell cycle classification of hiPSC subpopulations

To test the hypothesis that the subpopulations classified as core pluripotent (subpopulation one) or proliferative (subpopulation two) would harbor specific cell cycle signatures, we predicted the cell cycle phase of each cell (Scialdone et al., 2015; Lun et al., 2016). To estimate phase scores for each cell, the prediction method uses gene expression data and a reference training set containing prior ranks of relative expression of “marker pairs”, in which the sign of each pair changes between cell cycle phases (Scialdone et al., 2015; Leng et al., 2015) (**Figure 4f**). In particular, we observed a significantly higher percentage of cells in S (synthesis) phase belonged to subpopulation two (14.2%) compared to subpopulation one (8.4%: Fisher’s exact test, *P* < 2.2 × 10^−16^), and a significantly higher percentage of cells in G1 (50.3%) belonged to subpopulation one compared to subpopulation two (45.6%: Fisher’s exact test, *P* < 2.2 × 10^−16^). **Table 2** summarizes the proportions of cells in each cell cycle phase grouped by subpopulation. These data confirm our previous findings that subpopulation two is more proliferative and likely forms the reversible, self-renewing component of the pluripotent population as a whole.

**Table 2.** Prediction of cell cycle phases for each of the 18,787 cells in four subpopulations.

| Subpopulation ID | Phase | Cell Number | Cluster Percent | Cluster Total | Phase Percent | Phase Total |
| --- | --- | --- | --- | --- | --- | --- |
| 1 | G1 | 8084 | **89 (\*\*\*)** | 9083 | 50.3 | 16075 |
| 2 | G1 | 7324 | 81.6 | 8977 | 45.6 | 16075 |
| 3 | G1 | 482 | 91.6 | 526 | 3 | 16075 |
| 4 | G1 | 185 | 92 | 201 | 1.2 | 16075 |
| 1 | G2M | 239 | 2.6 | 9083 | 37.3 | 640 |
| 2 | G2M | 374 | 4.2 | 8977 | 58.4 | 640 |
| 3 | G2M | 21 | 4 | 526 | 3.3 | 640 |
| 4 | G2M | 6 | 3 | 201 | 0.9 | 640 |
| 1 | S | 760 | 8.4 | 9083 | 36.7 | 2072 |
| 2 | S | 1279 | **14.2 (\*\*\*)** | 8977 | 61.7 | 2072 |
| 3 | S | 23 | 4.4 | 526 | 1.1 | 2072 |
| 4 | S | 10 | 5 | 201 | 0.5 | 2072 |

\*\*\* Cluster 1 vs 2: $P < 2.2 \times 10^{-16}$ , Fisher's Exact Test

## Discussion

While methods to dissect cell subpopulations at single cell resolution such as FACS and immunohistochemistry have been available, a comprehensive profiling of transcriptional state(s) defining functionally distinct cell subpopulations comprising a ‘homogenous’ hiPSC cell line has not been described (Wilson et al., 2015; Kalkan et al., 2017). Although the heterogeneity of pluripotency states in an induced pluripotent cell culture is widely recognized, the quantitative characterization of subpopulations, identification of markers for subpopulations and prediction of transition potential among them have not been investigated. To address this, we generated and analyzed the largest hiPSC single-cell transcriptomics dataset to date, from five biological replicates of an engineered WTC-CRISPRi hiPSC line (Mandegar et al., 2016). The 18,787 high-quality transcriptomes that comprise this hiPSC population, collectively expressing 16,064 genes, provided strong statistical power for unbiased computational decomposition of cellular and transcriptional heterogeneity.

We developed a high-resolution and deterministic clustering method (UHRC) that does not require a predetermined cluster number. Across five separate biological replicates, we consistently found the existence of two main subpopulations including a core pluripotent and a pluripotent-proliferative subpopulation, accounting for 96.1% of all cells profiled. Comparison of transcriptomes between subpopulations revealed a cell proliferation gene network, coordinately regulated by two transcription factors *SALL4, ZIC1* and the *NR5A1* signaling receptor, together with the well-characterized pluripotency regulators *POU5F1, SOX2*, and *NANOG*. Recently, Kalkan *et al* (2017) detected higher expression of *Sall4* and *Nr5a2* by single-cell RNA-seq in the early undifferentiated state (2i) of mouse ESCs compared to later, more differentiated time-points. The separation of two major subpopulations on the basis of cell proliferation states may in part be explained by evidence that reprogramming is commonly a stochastic process dependent on cell-proliferation rate (Hanna et al., 2009). It remains to be determined whether these subpopulations generally reflect a common feature of pluripotency in hESC or hiPSC populations. Furthermore, the high sensitivity of the UHRC method allowed the detection of two smaller subpopulations (2.8% and 1.1% of the total cells) with transcriptional signatures of pluripotency but primed for differentiation based on enriched signaling pathways and gene ontologies related to lineage specification. Interestingly, from analysis of expression heterogeneity within and between subpopulations, we found higher variability in these two subpopulations compared to the remaining cells. This observation is consistent with recent single-cell studies showing that the transition from pluripotency to lineage commitment phase is characterized by high gene expression variability (Semrau et al., 2016) and by the gradual destabilization of the pluripotent stem cell networks (Bargaje et al., 2017).

Moreover, we developed a method to find novel gene markers and models capable of distinguishing these subpopulations. Our marker identification method enables objective selection of genes from a large set containing hundreds to thousands of differentially expressed genes. The novel markers outperform known markers in predicting subpopulations. We developed a machine learning approach that can be widely applied to optimize prediction models based on single-cell transcriptomics data to classify cells into subpopulations at a high accuracy. Identifying cell types is often based on immunostaining, FACS, or targeted PCR quantification of a small number of markers (Tsankov et al., 2015; Kalkan et al., 2017). Here, we constructed an unbiased classification model based on differential gene expression selected by the LASSO penalized maximum likelihood optimization procedure. The procedure does not require model tuning by selecting parameters based on prior knowledge. We identified large, previously unreported groups of genes as new predictors for pluripotency and showed that prediction models from differentially expressed genes performed better than models built from known markers. Our model is especially effective at identifying genes and estimating their effects in the prediction of complex phenotypes (polygenic traits), which are controlled by multiple genetic loci and where each locus has a small individual effect (Yang et al., 2010a; Boyle et al., 2017). Our results support the use of an unbiased and transcriptome-wide approach to developing gene prediction models, which can leverage subtle expression changes in a large number of genes to accurately classify cell types (subpopulations) and discover novel gene markers for important phenotypes. The results of these predictions can generate important hypotheses around genes and networks to be validated by functional genomics assays.

Importantly, using LTS, our quantitative and directional prediction method, we observed that these multiple iPSC subpopulations have the potential to transition between pluripotency states, which was further supported by two independent analysis methods, Monocle2 and diffusion pseudotime. The results of these trajectory analyses revealed the dynamic transition among subpopulations that ensures both cell state-renewal (transition between subpopulation two and subpopulation one) and differentiation capacity (subpopulation one to subpopulations three and four). This is especially the case for subpopulations 1 and 2, which are on a continuum yet are distinguishable from one another based on their transcriptional profiles. However, between subpopulations 2 to 3 and 2 to 4, the continuum is less clear, suggesting greater distinction between each subpopulation (**Figure 4**). Prior work using microarray gene expression arrays on human embryonic stem cells (hESCs) suggested a continuum of pluripotent states exists (Hough et al., 2009). Our finding is consistent with a study on early developmental progression from naive pluripotency in human epiblast and embryonic stem cells to more differentiated tissues such as trophectoderm and primitive endoderm (Yan et al., 2013). Using single cells, the authors reported a strong reduction in the expression of pluripotency markers during the cell transition from pluripotent to differentiated. A second study investigating early developmental progression from naïve pluripotency in mouse ESCs (Kalkan et al., 2017) showed reduced expression of pluripotency markers occurred prior to the activation of lineage specification markers.

Despite the large number of cells sequenced, this study was limited in that only 3’ mRNA was sequenced, and thus there remained variation between cell populations that could not be detected. In addition, our observations were obtained from one iPSC line and may not reflect the general behavior of all pluripotent stem cells. Nevertheless, our aim was to deconvolute a ‘homogeneous’ hiPSC population, and inclusion of transcriptional sequence data from other RNA species in the future will likely improve our ability to further delineate subpopulations of cells. Furthermore, we confirmed that the genes selected were expressed in 71 HipSci datasets (Streeter et al., 2017), and that relative expression level among genes was consistent between scRNA and bulk RNA sequencing. The parental cell line selected for this study, WTC-CRISPRi hiPSCs (Mandegar et al., 2016), is an important system for targeted transcription inhibition, and is a key feature for functional genomics studies that build on this dataset to study the biology of pluripotency. Coupled with high-throughput single-cell RNA-seq, our innovative computational methods have revealed the intrinsic characteristics that distinguish subpopulations of pluripotent stem cells. Future work is required to expand this analysis to multiple hiPSC and hESC lines to identify common features of single-cell subpopulations in pluripotency.

## Methods

### Cell culture

Undifferentiated human induced pluripotent stem cells (hiPSC; WTC-wild type C) were provided courtesy of Bruce Conklin (UCSF and Gladstone Institutes) as previously described (Mandegar et al., 2016). Cells were maintained on Vitronectin XF (STEMCELL Technologies, cat. no. 07180) and cultured in mTeSR1 (STEMCELL Technologies, cat. no. 05850). Cytogenetic analysis by Giemsa banding showed a normal 46, XY male karyotype. For scRNA-seq, samples one and two were harvested from a single plate using Versene, split into two technical replicates resuspended in Dulbecco’s PBS (dPBS; Life Technologies, cat. no. 14190-144) with 0.04% bovine serum albumin (Sigma, cat. no. A9418-50G) and immediately transported for cell sorting. For samples 3-5 cells were harvested from individual plates using 0.25% Trypsin (Life Technologies, cat. no. 15090-046) in Versene, neutralized using 50% fetal bovine serum (HyClone, cat. no. SH30396.03) in DMEM/F12 (Life Technologies, cat. no. 11320-033), centrifuged at 1200 rpm for 5 minutes and re-suspended in dPBS + 0.04% BSA.

### Cell sorting

Viable cells were sorted on a BD Influx cell sorter (Becton-Dickinson) using Propidium Iodide into Dulbecco’s PBS + 0.04% bovine serum albumin and retained on ice. Sorted cells were counted and assessed for viability with Trypan Blue using a Countess automated counter (Invitrogen), and then resuspended at a concentration of 800-1000 cells/μL (8 × 10^5^−1 × 10^6^ cells/mL). Final cell viability estimates ranged between 80-93%.

### Generation of single cell GEMs and sequencing libraries

Single cell suspensions were loaded onto 10X Genomics Single Cell 3’ Chips along with the reverse transcription (RT) mastermix as per the manufacturer’s protocol for the Chromium Single Cell 3’ Library (10X Genomics; PN-120233), to generate single cell gel beads in emulsion (GEMs). Reverse transcription was performed using a C1000 Touch Thermal Cycler with a Deep Well Reaction Module (Bio-Rad) as follows: 55°C for 2h; 85°C for 5min; hold 4°C. cDNA was recovered and purified with DynaBeads MyOne Silane Beads (Thermo Fisher Scientific; Cat# 37002D) and SPRIselect beads (Beckman Coulter; Cat# B23318). Purified cDNA was amplified as follows: 98°C for 3min; 12×(98°C for 15s, 67°C for 20s, 72°C for 60s); 72°C for 60s; hold 4°C. Amplified cDNA was purified using SPRIselect beads and sheared to approximately 200bp with a Covaris S2 instrument (Covaris) using the manufacturer’s recommended parameters. Sequencing libraries were generated with unique sample indices (SI) for each sample. Libraries for samples 1-3 and 4-5 were multiplexed respectively, and sequenced on an Illumina NextSeq 500 (NextSeq control software v2.0.2/ Real Time Analysis v2.4.11) using a 150 cycle NextSeq 500/550 High Output Reagent Kit v2 (Illumina, FC-404-2002) in stand-alone mode as follows: 98bp (Read 1), 14bp (I7 Index), 8bp (I5 Index), and 10bp (Read 2).

### Bioinformatics mapping of reads to original transcripts and cells

Processing of the sequencing data into transcript count tables was performed using the Cell Ranger Single Cell Software Suite 1.2.0 by 10X Genomics (http://10xgenomics.com/). Raw base call files from the NextSeq 500 sequencer were demultiplexed, using the *cellranger mkfastq* pipeline, into sample-specific FASTQ files. These FASTQ files were then processed with the *cellranger count* pipeline where each sample was processed independently. First, *cellranger count* used STAR (Dobin et al., 2013) to align cDNA reads to the hg19 human reference transcriptome, which accompanied the Cell Ranger Single Cell Software Suite 1.2.0. We note that, since the expression data is limited to the 3’ end of a gene and we used gene-level annotations, differences between reference versions, such as GRCh38, are unlikely to significantly alter conclusions. Aligned reads were filtered for valid cell barcodes and unique molecular identifiers (UMI) and observed cell barcodes were retained if they were 1-Hamming-distance away from an entry in a whitelist of known barcodes. UMIs were retained if they were not homopolymers and had a quality score > 10 (90% base accuracy). *cellranger count* corrected mismatched barcodes if the base mismatch was due to sequencing error, determined by the quality of the mismatched base pair and the overall distribution of barcode counts. A UMI was corrected to another, more prolific UMI if it was 1-Hamming-distance away and it shared the same cell barcode and gene. *cellranger count* examined the distribution of UMI counts for each unique cell barcode in the sample and selected cell barcodes with UMI counts that fell within the 99^th^ percentile of the range defined by the estimated cell count value. The default estimated cell count value of 3000 was used for this experiment. Counts that fell within an order of magnitude of the 99^th^ percentile were also retained. The resulting analysis files for each sample were then aggregated using the *cellranger aggr* pipeline, which performed a between-sample normalization step and merged all 5 samples into one. Post-aggregation, the count data was processed and analyzed using a comprehensive pipeline assembled and optimized in-house as described below.

### Preprocessing

To preprocess the mapped data, we constructed a cell quality matrix based on the following data types: library size (total mapped reads), total number of genes detected, percent of reads mapped to mitochondrial genes, and percent of reads mapped to ribosomal genes. Cells that had any of the 4 parameter measurements higher than 3x median absolute deviation (MAD) of all cells were considered outliers and removed from subsequent analysis (**Supplemental Table S2**). In addition, we applied two thresholds to remove cells with mitochondrial reads above 20% or ribosomal reads above 50% (**Supplemental Table S2**). To exclude genes that were potentially detected from random noise, we removed genes that were detected in fewer than 1% of all cells. Before normalization, abundantly expressed ribosomal protein genes and mitochondrial genes were discarded to minimize the influence of those genes in driving clustering and differential expression analysis.

### Data normalization

Two levels of normalization were performed to reduce possible systematic bias between samples and between cells. To reduce potential confounding effects caused by differences in sequencing depths between five samples, a subsampling process (Zheng et al., 2017) was used to scale the mean mapped reads (MMR) per cell of all samples down to the level of the sample with the lowest MMR. For each sample, a binomial sampling process randomly selected reads and UMIs for each gene in a cell at a sample-specific subsampling rate. The subsampling rate for each sample was determined using the ratios of expected total reads (given the expected mean reads per cell (minimum MMR of all samples), the known number of cells, and the fraction of mapped reads to total reads) to the original total mapped reads (equation 1). Following resampling, the MMRs for the five samples were scaled, while the expression data distribution for genes in all cells of the sample was for genes in all cells of the sample was maintained.

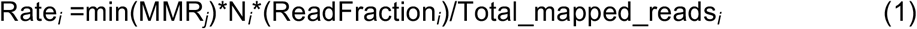

where: min(MMR_*j*_) is the minimum MMR of all samples to be merged; N_*j*_ is the number of cells in sample *j*; ReadFraction_*i*_ is the ratio of confidently mapped reads in a cell to the total number of reads detected for that cell in sample *i*; Total_mapped_reads_i_ is the total number reads that share the same cell barcode. For each gene in each cell, we performed a random binomial sampling process of reads with the probability equal the subsample rate calculated in equation (1). This process is more robust than standard scaling options because it takes into account unique read information associated with mapped genes and cells.

To reduce cell-specific bias, possibly caused by technical variation such as cDNA synthesis, PCR amplification efficiency and sequencing depth for each cell, expression values for all genes in a cell were scaled based on an estimated cell-specific size factor. Before normalization, counts were log_2_-transformed (by log_2_(count+1)) to stabilize variance due to the large range of count values (spanning 6 orders of magnitude, **Supplemental Figure S1e**). To estimate the scaling size factor for each cell, a deconvolution method (Lun et al., 2016) was applied for summation of gene expression in groups of cells. This summation approach reduced the number of stochastic zero expression of genes that are lowly expressed (higher dropout rates), or genes that are turned on/off in different subpopulations of cells.

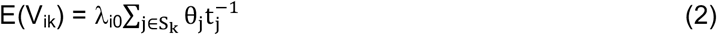

Where S_k_ is a pool of cells, V_ik_ is the sum of adjusted expression value (Z_ij_ = θj\**λ*_*i*0_, where *λ*_*i*0_ is the expected transcript count and θ_j_ is the cell-specific bias) across all cells in pool V_k_ for gene *i*, 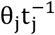 is the cell-specific scale factor for cell *j* (where *t*_j_ is the constant adjustment factor for cell *j*).

The estimated size factor of a gene in in pool S_k_, named as E(R_ik_), is the ratio between the estimated V_ik_ and the average Z_ij_ across all cells in the population. 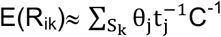 where 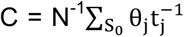, where *N* is the number of cells, S_0_ represents all cells and is a constant for the whole population and thus can be set to unity or ignored. The cell pools were sampled using a sliding window on a list of cells ranked by library size for each cell. Four sliding windows with 20, 40, 60, and 80 cells were independently applied and results were combined to generate a linear system that can be decomposed by QR decomposition to estimate 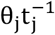 size factor for each of the cell. The normalized counts are the results of taking the raw counts divided by cell-specific normalized size factors.

### Developing a publically accessible data resource

To make this valuable single-cell human pluripotent single-cell dataset publicly accessible we created an interactive R shiny server at http://computationalgenomics.com.au/shiny/hipsc/. The server contains user-friendly data exploration and representation tools. Users can interactively explore the expression of any selected gene from the 16,064 genes in each cell of the 18,787 cells. The data specific for each subpopulation is also calculated. For each gene, a summary table, tSNE plot and a density plot can be generated and the results are downloadable.

### Analyzing transcriptional heterogeneity in a population of single cells

To assess transcriptional heterogeneity among cells and genes, we first removed potential variation due to technical sources by the subsampling process and the cell-specific normalization as described above. Depending on experimental design, an additional step using a generalized linear model (GLM) to regress out other potential confounding factors can be included. After reducing technical variation via normalization, we calculated the coefficient of variation and expression dispersion of each gene across all cells. For cell-to-cell variation, we first performed principal component analysis (PCA) on general cell data, which included percent counts of the top 100 genes, total number of genes, percent of mitochondrial and ribosomal genes. To investigate variation between genes, the distribution of dispersion across a range of expression values was calculated (equation 3). This approach is useful because technical variation often appears greater in lowly expressed genes than in more abundant genes (Shalek et al., 2013). Denoting x_i_ as the vector of expression values (in counts-per-million) for gene *i* across all cells, we use the following formula to compute coefficient of variation (*cv*):

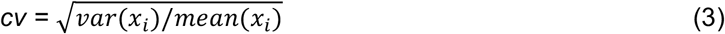

We estimated the BCV (biological coefficient of variation) with an empirical Bayesian approach to estimate dispersion between genes and between samples (McCarthy et al., 2012). Common dispersion (shared dispersion value of all genes), trended dispersion (mean dispersion trend for lowly expressed genes to abundant genes), and gene-specific dispersion was estimated to reflect variation of all genes across the whole population (**Supplemental Figure S9**).

### Dimensionality reduction

After merging five samples, preprocessing, and normalizing the dataset was scaled to *z*-distribution and PCA was performed for dimensionality reduction using the prcomp function in R (McCarthy et al., 2016). To assess PCA results, we examined the top genes that were most correlated to PC1 and PC2, and the distribution of cells and percent variance explained by the top five PCs. Importantly, the optimal number of PCs explaining the most variance in the dataset was determined using a Scree test calculated by the fa.parallel function in the psych package. The fa.parallel was run based on expression data for the top variable genes.

Cells are represented using *t*-SNE and diffusion map (van der Maaten and Hinton, 2008). We used the Rtsne package v0.1.3 with the normalized expression data (16,064 genes by 18,787 cells) to calculate a three-dimensional *t*-SNE projection dataset (16,064 genes by three *t*-SNE dimensions), which was then combined with other data types to display cells on two- and three-dimensional *t*-SNE plots.

### Clustering

We developed an unsupervised clustering method that does not require the number of clusters to be specified prior to the analysis so that small and big clusters can be determined automatically. We first computed a cell-PCA eigenvector matrix based on the full expression dataset (all 18,787 cells and over 15,953 genes) to reduce the full dataset to a 18,787 cells × 10 first orthogonal principal components (PCs). The first 10 PCs that explained for most of the variance of the full expression matrix were used for computing cell-to-cell Euclidean distance. We applied an agglomerative hierarchical clustering (HAC) procedure using Ward’s minimum variance (minimal increase of sum of squared) method aiming at finding compact clusters to minimize total within-cluster variance. By using Ward’s linkage method, cells and branches are joined iteratively from the bottom (each cell is one branch) to the top (all cells form one cluster), resulting in a complete dendrogram tree. To consider the broadest possible solution space for finding clusters from the tree without a constraining threshold, we allowed a cut height of 99% of the range between the 5th percentile and the maximum of the joining height on the dendrogram, allowing the entire set of branches in the original tree to be considered in the cluster search space.

To determine the number of subpopulations, branches of the dendrogram were pruned by a Dynamic Tree Cut method, which uses an approach that does not attempt to merge all branches with distance lower than a constant (supervised) height cutoff (Langfelder et al., 2008). The dynamic merging process is a hybrid method that uses the dendrogram tree shape to define the bottom level clusters (based on number of cells, distance between cells, core of the branches and gaps between branches) and uses the dissimilarity measure to merge cells/branches (with lowest dissimilarity) into the initially defined clusters. The dynamic, iterative decomposition and combination of clusters enables the sensitive detection of outliers and nested clusters. To ascertain the clustering results are stable. We performed cluster stability analysis and resolution analysis. We narrowed the clustering search space by running 10 clustering iterations at the bottom 25% height of the entire tree. In each iteration, we decreased the pruning window by 2.5%. The bottom 25% of the tree was used in the stability analysis to keep clusters at the highest resolution, allowing detection of small clusters. Among all 10 independent clustering iterations, we consistently found 4 clusters (**Supplemental Figure S4**).

To allow for the detection of the smallest subpopulation, we considered any cluster with more than 20 cells to be significant in size, thereby enabling the detection of a cluster as small as 0.1% of the total cells. With these two parameters, the whole tree and the smallest clusters are considered without predetermined parameters.

Cluster information was then overlaid on cells in 2-3-dimensional t-SNE plots. Notably, the clusters were the results from HAC analysis and were not dependent on the t-SNE plots, which were used purely for representation purposes.

### Cell cycle analysis

To assess whether the clustering assignments were affected by the differences in cell cycle phases, we applied a machine learning prediction model to predict cell cycle phase (Scialdone et al., 2015; Lun et al., 2016). The model uses scRNA gene expression data and a reference training set (prior-knowledge) on relative expression of “marker pairs”, in which the sign of each pair changes between cell cycle phases (Scialdone et al., 2015). Scores for each of the three phases G1, G2M and S were estimated based on the proportion of training pairs having the sign changed in each phase relative to the other phases. The human training set was from Leng et al. (2015).

### Differential expression analysis

To select genes that distinguish subpopulations, we performed pairwise differential expression analysis between cells in pairs or groups of clusters by fitting a general linear model and using a negative binomial test as described in the DESeq package (Anders and Huber, 2010). Each cell was considered as one biological replicate in each cluster. We found that the shrinkage estimation of dispersion approach used in DESeq produced stable estimation of scale factors for genes and cells between clusters and was more conservative in detecting differentially expressed genes, especially when comparing subpopulations with a larger number of cells, such as subpopulation one and two, to subpopulations with small cell numbers, such as three and four. Specifically, DESeq detected fewer DE genes that expressed highly in a small proportion of cells in a subpopulation while remaining cells in that subpopulation had zero or very low expression. Significantly differentiated genes were those with Bonferroni threshold *P*-values less than 5% (*p* < 3.1 × 10^−7^).

### Subpopulation Classification Analysis

To develop predictive models based on single cell transcriptomics data, we applied the Least Absolute Shrinkage and Selection Operator (LASSO) procedure. LASSO selects gene predictors for classifying cells into one of the four subpopulations. Briefly, penalized logistic regression was applied to fit a predictor matrix (training set) containing expression values of the top differentially expressed genes in 50% of the total cells, or for the smallest subpopulation four, a subsample of a randomly selected 10% of the total cells, and a response variable vector assigning cells into one of the subpopulations (dichotomous variable, with two classes: belonging to the subpopulation or belonging to the remaining cells). The LASSO procedure optimizes the combination set of coefficients for all predictors in a way that the residual sum of squares is smallest for a given lambda value (Friedman et al., 2010).

We applied the glmnet R package to estimate parameters by a penalized maximum likelihood procedure as shown in the equation (4) so that insignificant genes *j* (not explaining variance) were removed (coefficients shrunken to 0):

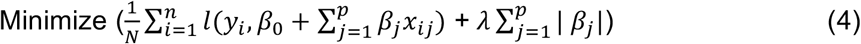

where **x**_i_ = (x_*i*1_, x_*i*2_, …, x_*i*p_) is a vector of expression values of *p* genes in cell C_*i*_; *y_i_* is the cell subpopulation class of the cell C_*i*_; 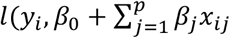 is the negative log-likelihood for C_*i*_; and *λ* is a tuning parameter that controls the shrinkage penalty of the coefficients.

For each training dataset, an optimal *λ* and a set of gene predictors can be determined by a 10-fold cross-validation procedure to select the *λ* that produced the minimum classification errors. The LASSO procedure optimizes the combination set of coefficients for all predictors in a way that the residual sum of squares is smallest for a given lambda value [38]. In other words, the LASSO procedure identified an optimal combination of genes (predictors) and fitted a logistic regression model. The logistic regression model is a generalized linear model in which the response variable belong to the binomial family, in which expression values of the selected genes were predictors and the binary labels (classes) of cells were response variables. The fitted model could either explain the highest deviance, compared to the full model, or classify cells to subpopulations with the lowest 10-fold classification error. The glmnet R package was applied to select top genes from the differentially expressed genes that contributed to classifying cells into each subpopulation (Tibshirani et al., 2012). To calculate the model prediction accuracy, we trained the LASSO model using one subsampled dataset and then evaluated the trained model in predicting a new, non-overlapping subsampled dataset. By comparing the prediction results with the known subpopulation label of each cell, the model prediction accuracy was calculated. We applied a Bootstrap procedure to calculate classification accuracy for 100 iterations.

### Pseudotime analysis

We applied the optimal LASSO classification model trained in the procedure above to estimate transition scores between subpopulation. For two subpopulations, the transition score is the percentage of cells in the target subpopulation that are classified as belonging to the original subpopulation and not belonging to the other class. For each subpopulation, the logistic regression model (with LASSO selected genes and corresponding coefficients) was fitted into a new expression dataset for all cells in another subpopulation. The model estimates the conditional class probability of a cell C_*i*_ belonging to a class k (0 or 1 ≡ belong or not belong to) given the gene expression profile **x_*i*_** of the *p* LASSO-selected genes **x_*i*_**:

The conditional class probabilities of cell C_*i*_ belonging to class *k* is then the linear combination of selected genes, and can be estimated as:

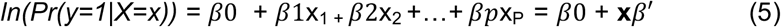

where *βj* is a coefficient for gene *j* (*βj*=0 if the gene *j* is not a predictor of the class). The coefficient vector *β*= (*β*0,*β*1….*βp*) can be calculated by maximum likelihood estimation and the deviance that best explains the variance compared to the full model, as in equation (4). The predicted probability of a cell C_*i*_ being in a subpopulation 1 or 0 is estimated by replacing *β* and gene expression values in the regression equation (5).

Results from our novel trajectory analysis method was compared to the two state-of-the-art pseudotime analysis approaches, namely Diffusion pseudotime (Haghverdi et al., 2016) and Monocle2 (Qiu et al, 2017). The Diffusion method, implemented in Destiny (version 2.0.8), applies a diffusion-like random walk algorithm to estimate the cell’s probabilities, based on a weighted nearest neighbor graph, of transitioning into another cell at a different fate/state, thereby inferring a differentiation trajectory. The diffusion approach is not dependent on dimension reduction. The Monocle2 package (version 2.2.0) applies an unsupervised manifold learning technique, namely reverse graph embedding (DDRTree - Discriminative Dimensionality Reduction for learning principal graphs), to learn an optimal path (a curved manifold in a low dimensional space) that approximates the structure of high-dimensional data and reversely map cells on this path back to the original multidimensional space so that nearby cells on the manifold are also nearby cells on the original space (Qiu et al, 2017). Both Monocle2 and Diffusion methods estimate pseudotime, which is a measure of how far a cell progresses in the differentiation process compared to a root cell. Pseudotime is therefore the distance between two cells in a modelled trajectory and is estimated independently from the experimental time.

### Pathway and gene functional analysis

To functionally characterize the four subpopulations, we performed a network analysis using significant DE genes between cells within a subpopulation and the remaining cells, or between cells in pairs of subpopulations. We used Cytoscape to apply three main programs: GeneMANIA (Warde-Farley et al., 2010), with a comprehensive background database containing 269 networks and 14.3 million interactions, the Reactome functional interaction network analysis, a reliably curated protein functional network (Wu et al., 2010), and the STRING protein-protein interaction database (Szklarczyk et al., 2015).

## Data access

The raw and processed data is available at ArrayExpress under E-MTAB-6687. All code for figures and analysis are on GitHub (https://github.com/IMB-Computational-Genomics-Lab/hiPSC_paper_2018) and available as Supplemental_Code.zip.

## Acknowledgments

Sequencing was performed by the Institute for Molecular Bioscience Sequencing Facility at the University of Queensland. The authors thank Stacey Andersen for valuable discussions and Liam Crowhurst for contributing to the development of the interactive Shiny apps. This work was supported by the Australian National Health and Medical Research Council (NHMRC) grant APP1083405 and APP1107599.

## Author contributions

J.E.P. designed the study, acquired funding and led analysis. N.J.P. led the cell culture and informed on the interpretation of results. S.W.L., H.S.C., T.J.C.B. and A.N.C. performed molecular experiments. Q.H.N, S.W.L., A.S performed analysis. All authors wrote and edited the manuscript.

## Disclosure Declaration

The authors declare there are no conflicts of interest.

